# Uncertainty-Aware Model Selection with a Calibrated Probability-Generating-Function-Based Bayesian Information Criterion

**DOI:** 10.64898/2026.09.11.750968

**Authors:** Yiling Wang, Zhanpeng Shu, Furong Gao, Edward Z. Cao

**Author notes:** Corresponding author Edward Z. Cao.

## Abstract

Selecting stochastic gene-expression models from single-cell counts requires balancing goodness of fit against unnecessary mechanistic complexity. The probability-generating-function-based Bayesian information criterion (PGF-BIC) combines covariance-weighted fitting in generating-function space with a complexity penalty, allowing candidate models to be compared without reconstructing their full count distributions. However, its conventional zero-threshold rule does not account for sampling uncertainty in the fitted score difference and may therefore favor overly complex models in finite samples. To address this limitation, we develop an uncertainty-aware PGF-BIC rule that selects the more complex model only when its score advantage exceeds a data-driven threshold. We use Cantelli’s one-sided inequality to motivate a selection margin expressed in terms of a standard deviation. To determine this scale, we use influence functions to quantify sensitivity to small perturbations in the data distribution and obtain a first-order description of sampling fluctuations. The resulting variance estimate accounts for variability in both the empirical probability generating function and the estimated covariance weights, yielding a data-driven threshold for assessing the complex model’s score advantage. A Poisson versus Bursty benchmark shows that the calibrated rule reduces incorrect selection of the more complex model. The calibration requires neither resampling nor additional optimization, incorporating sampling uncertainty into model selection while retaining the computational efficiency of PGF-BIC.

## I. Introduction

Stochastic gene expression can generate substantial variation in mRNA abundance among genetically identical cells under similar environmental conditions [1], [2]. Stochastic models relate this variability to promoter switching, transcript synthesis and degradation [3], with kinetic parameters determining the predicted count distributions. Inferring these parameters from single-cell counts thus provides a quantitative description of gene-expression dynamics [4], [5].

Parameter inference involves repeated comparisons between model predictions and observations, so the representation of count distributions directly affects computational efficiency. Maximum likelihood estimation uses count probabilities, but their computation may require costly numerical solutions of the chemical master equation over large state spaces [6]. Moment-based methods instead match means, variances or higher-order moments [7]. Higher-order sample moments, however, are particularly sensitive to sampling noise, which can reduce the reliability of parameter estimates when cell numbers are limited [5]. Probability-generating-function (PGF) methods offer an alternative by fitting model PGFs directly to empirical PGFs computed from the data [4], [8], avoiding the computation of count probabilities. Published studies have shown that PGF-based inference can achieve parameter-estimation accuracy comparable to maximum likelihood estimation with substantially shorter computation times [4], [5].

Efficient parameter inference makes it possible to fit candidate models to the same count data and compare their predictions. Yet different mechanisms can yield similar count distributions, even when their parameters describe distinct biological processes [9], [10]. Parameter fitting must therefore be complemented by model selection to assess whether the data support additional mechanistic complexity. Common approaches include cross-validation, the Akaike information criterion and the Bayesian information criterion (BIC) [11], [12]. Existing PGF workflows use cross-validation to assess fit on held-out observations [4], [5], but repeated fitting increases the cost of analyzing many genes and candidate models. Given a suitable likelihood, BIC offers a less computationally demanding comparison by combining each candidate’s full-data fit with a complexity penalty.

To enable such a comparison in PGF space, Wang et al. constructed a Gaussian quasi-likelihood that accounts for the sampling covariance of empirical PGF values [13]. This Gaussian approximation describes the sampling distribution of the empirical PGF, not the distribution of the underlying RNA counts. The resulting criterion supports covariance-weighted parameter estimation and, after adding a complexity penalty, yields PGF-BIC.

Computational efficiency alone does not resolve how small score differences should be interpreted. Standard PGF-BIC selects the more complex model whenever its penalized score is lower. However, both the empirical PGF and the covariance weights are estimated from the same cells, so the fitted score difference fluctuates across samples. When candidate distributions nearly coincide, these fluctuations can make a reproducible fitting advantage difficult to distinguish from an apparent improvement caused by sampling noise. Covariance weighting determines how discrepancies across PGF nodes contribute to the score, while the BIC penalty imposes a cost for additional parameters. Together, they define the comparison score but leave a further question: how large must the complex model’s score advantage be, relative to its sampling variability, to justify selecting that model?

To reduce incorrect selection in finite samples, we propose an uncertainty-aware PGF-BIC rule that selects the more complex model only when its score advantage exceeds a data-driven threshold. Cantelli’s one-sided inequality [14] motivates a threshold expressed in terms of a standard deviation. To estimate this scale, influence functions [15], [16] link sensitivity to small distributional perturbations with first-order sampling fluctuations, accounting jointly for variability in the empirical PGF and covariance weights estimated from the same cells. A Poisson versus Bursty benchmark demonstrates reduced incorrect selection of the more complex model. The calibration preserves the original fits and complexity penalty and requires neither resampling nor additional optimization.

## II. PGF-Based Model Comparison AND A Motivating Example

### A. Count Data and PGF Representation

We compare model predictions with observed counts in PGF space [13]. Evaluating the model PGF and the empirical PGF at the same fixed points provides a common vector representation for fitting and model comparison.

Let 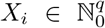 denote the count vector for cell *i*. We assume that observations from *n* cells measured under a common condition satisfy 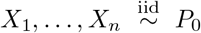, where the data-generating distribution *P*_0_ is unknown. We focus on selecting between a simpler model *M*_1_ and a more complex model *M*_2_, with parameters *θ*_1_ ∈ Θ_1_ ⊆ ℝ^*p*_1_^ and *θ*_2_ ∈ Θ_2_ ⊆ ℝ^*p*_2_^, respectively, where *p*_2_ > *p*_1_. Both models are fitted to and compared on the same observations, without prespecifying either as the data-generating mechanism. We assume that both model PGFs can be evaluated directly at the chosen points.

For a count distribution *P*, ⟨*f* (*X*)⟩_*P*_ denotes the expectation of *f* (*X*) under *P*; we omit the subscript when the distribution is clear. The PGF is defined as [17]

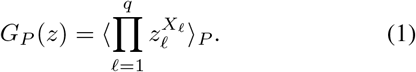

For each *z* ∈ (0, 1)^*q*^, the PGF is the expectation of a bounded transformation of the counts and can therefore be estimated by averaging the transformed observations. Evaluating it at *m* nodes *z*_1_, …, *z*_*m*_ (0, 1)^*q*^, fixed in advance and independently of the data, yields a finite-dimensional representation. Define 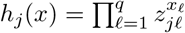 and *h*(*x*) = (*h*_1_(*x*), …, *h*_*m*_(*x*))^T^. The PGF vector for a distribution *P* is then *g*(*P*) = ⟨*h*(*X*)⟩_*P*_, with empirical estimate

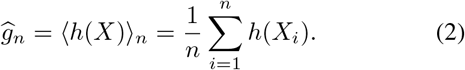

Here and below, the subscript *n* in ⟨·⟩_*n*_ denotes averaging over the observed cells. The empirical vector *ĝ*_*n*_ estimates *g*(*P*_0_), and the corresponding model vector has entries [*g*_*r*_(*θ*_*r*_)]_*j*_ = *G*_*r*_(*z*_*j*_; *θ*_*r*_). Evaluating both vectors at the same nodes allows parameter inference to proceed by adjusting *θ*_*r*_ to bring the model predictions closer to the empirical values, without reconstructing the full count distribution [8].

### B. Covariance-Weighted Fitting and PGF-BIC

With observations and predictions represented on a common grid, the next step is to quantify their discrepancy. We use covariance weighting to account for differences in estimation precision and correlations among empirical PGF components when measuring the fitting discrepancy [18]. Let Σ(*P*) = Cov_*P*_ {*h*(*X*)} denote the covariance of the feature vector under *P*. Using the centered observations *û u*_*i*_ = *h*(*X*_*i*_) − *ĝ*_*n*_, we estimate Σ(*P*_0_) and construct the fitting weights as

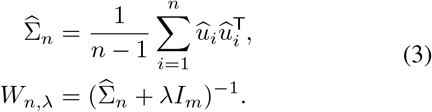

Here, *I*_*m*_ is the *m* × *m* identity matrix. Both candidates use the same fixed ridge parameter *λ ≥* 0 [19]; when 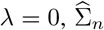 must be nonsingular. The corresponding population weight matrix is *W*_*λ*_(*P*) = {Σ(*P*) + *λI*_*m*_} ^−1^. For a fixed data-generating distribution *P*_0_, the population weights *W*_*λ*_(*P*_0_) are fixed, whereas their estimate *W*_*n,λ*_ varies with the observed cells.

For a reference count distribution *P*, define the covariance-weighted PGF distance for model *M*_*r*_ as *Q*_*r*_(*θ*_*r*_; *P*) = {*g*_*r*_(*θ*_*r*_) − *g*(*P*)}^T^*W*_*λ*_(*P*) {*g*_*r*_(*θ*_*r*_) − *g*(*P*)}. In practice, we fit each model to the same empirical PGF using common sample covariance weights that remain fixed during optimization [8], [13]. The parameter estimate is obtained by minimizing the corresponding sample distance:

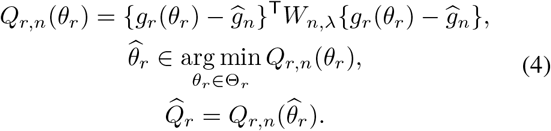

After fitting, we compare the two models using the following PGF-BIC scores and their difference [12], [13]:

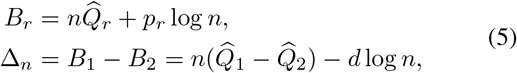

Here, *d* = *p*_2_ − *p*_1_ > 0, and *n* denotes the number of independent cells rather than PGF nodes. A positive Δ_*n*_ indicates a lower penalized score for *M*_2_, which the conventional rule selects; otherwise, it retains *M*_1_. This decision is based on quantities that vary across samples. Although the empirical PGF and covariance weights remain fixed during each optimization, both are re-estimated for a new dataset, and their joint variation affects the fitted score difference. The zero cutoff does not account for this uncertainty in Δ_*n*_, making small score advantages difficult to interpret when the fitted distributions are nearly indistinguishable.

### C. A Toy Example Illustrating Over-Selection by PGF-BIC

To examine this issue when the generating model is known, we simulate independent counts *X*_1_, …, *X*_*n*_ ~ Poisson(20) and compare a Poisson model (*M*_1_) with a Bursty model (*M*_2_) [10].

Let *M* denote mRNA and measure time in units of the mean transcript lifetime, so that each transcript degrades independently at unit rate. The two models are represented by

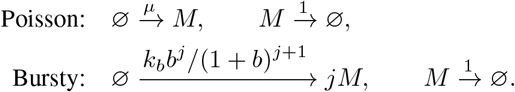

Here, *j* = 0, 1, … denotes the burst size, distinct from the sample size *n*. The Poisson model produces transcripts individually; in the Bursty model, burst events occur at rate *k*_*b*_ > 0 and independently produce geometrically distributed numbers of transcripts with mean *b* > 0 [20].

The corresponding stationary PGFs are

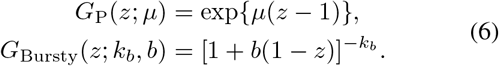

We estimate each model’s parameters separately: *µ* for the Poisson model and (*k*_*b*_, *b*) for the Bursty model, giving *p*_1_ = 1, *p*_2_ = 2 and *d* = 1. The stationary mean of the Bursty model is *k*_*b*_*b*; as *b →* 0 and *k*_*b*_*b → µ* > 0, its count distribution approaches Poisson(*µ*). This comparison therefore assesses whether the observed counts support dispersion beyond that described by the Poisson model.

We assess selection across sample sizes by generating 500 independent datasets for each *n* ∈ {100, 500, 1000}, with *n* cells per dataset. Both models are fitted on the common grid *z* = 0.50, 0.51, …, 0.99 with *λ* = 5 × 10^−16^. The correct-selection rates for Poisson are approximately 30%, 54.6% and 65.4% at *n* = 100, 500, 1000, respectively. Correct selection becomes more frequent as the sample size increases, but the conventional rule still selects the Bursty model in a substantial proportion of replicates.

Figure 1 compares the empirical count distribution with both fitted distributions for one dataset selected from the 500 independent replicates at each sample size. The fitted curves nearly overlap, yet some numerical comparisons give Δ_*n*_ > 0. Thus, the conventional rule can favor the Bursty model even when the data are generated by Poisson and the fitted distributions are almost indistinguishable.

**Fig. 1.**
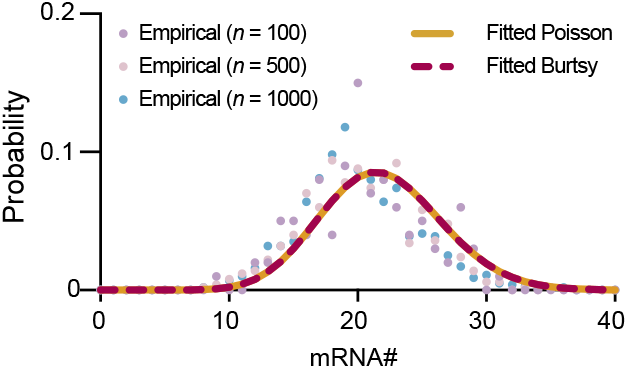
Empirical count distributions and Poisson and Bursty model fits, shown for one example dataset at each sample size.

Because Poisson is a boundary limit of the Bursty model, these examples illustrate finite-sample selection near a model boundary. The observed over-selection motivates a calibrated threshold that accounts for the sampling uncertainty of the fitted score difference when assessing whether the apparent advantage supports additional model complexity.

The relevant question is therefore not only which candidate has the lower score, but whether its advantage is sufficiently large relative to the sampling variability of the comparison. Once the sample size and candidate dimensions are specified, the penalty *d* log *n* is fixed, so this uncertainty arises from the difference between the minimized fitting distances. We retain the original fits and BIC penalty and replace the zero cutoff with a data-driven threshold, selecting the complex candidate *M*_2_ only when

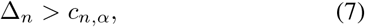

Here, *c*_*n,α*_ ≥ 0 and *α* ∈ (0, 1) is the target level for the probability of incorrectly selecting the more complex model. The next section first establishes the form of a one-sided uncertainty-based margin, then derives and estimates the fluctuation scale needed to compute it. The calibration uses the existing fits and accounts for joint variation in the empirical PGF and estimated covariance weights.

## III. Construction OF THE PGF-BIC Calibration Threshold

In the benchmark in Section II, the data are generated by the Poisson model, yet the conventional PGF-BIC rule selects the more complex Bursty model in a substantial proportion of replicates. To reduce such errors, we construct a selection threshold that accounts for sampling uncertainty in the score difference. We first formulate the calibration objective and use Cantelli’s inequality to determine the threshold’s form. We then derive and estimate the required sampling scale through influence functions, obtaining an explicit threshold and the associated model-selection rule.

### A. Calibration Objective and One-Sided Threshold Construction

Calibration adjusts only the decision threshold *c*_*n,α*_ in (7), leaving the fitted models and their PGF-BIC scores unchanged. To distinguish a population-level advantage from a sample-specific one, we use the criterion in Section II to define the difference between the models’ minimum distances to a reference distribution *P* :

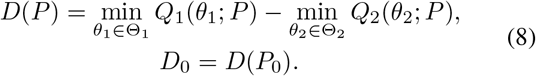

Let *θ*_0*r*_ minimize *Q*_*r*_(*θ*_*r*_; *P*_0_) for model *M*_*r*_. For a fixed *P*_0_, the population-optimal parameter 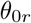 is fixed but generally unknown, whereas 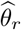 is fitted to the observed data using (4) and varies across samples.

To relate this population comparison to the observed score difference, let 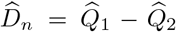 denote the fitted distance difference. Equation (5) gives 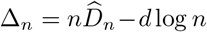. For fixed *n* and model dimensions, the penalty *d* log *n* is deterministic, so the sampling variability of Δ_*n*_ comes entirely from 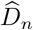.

We therefore examine 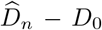, the deviation from the population reference.

The fitted distance difference 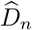 depends on the empirical PGF *ĝ*_*n*_ and empirical second moments 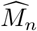. Write *g*_0_ = *g*(*P*_0_) and define the empirical and population second-moment matrices of *h*(*X*) as

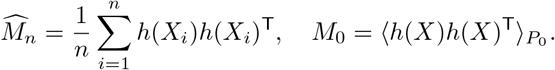

To characterize sampling fluctuations around *D*_0_, we treat 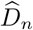 as a function of *ĝ*_*n*_ and 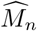 and take a first-order Taylor expansion at the population values (*g*_0_, *M*_0_), obtaining 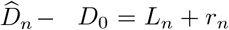. Here, *L*_*n*_ combines the linear contributions of the empirical PGF deviation *ĝ*_*n*_ − *g*_0_ and the empirical second-moment deviation 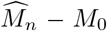 to the fitted distance difference and has zero population mean; *r*_*n*_ is the remainder. Derivations of the expansion and zero-mean property are given in Appendix A, (A.2)–(A.4). To account for the factor *n* in the PGF-BIC fitting term, set *Z*_*n*_ = *nL*_*n*_ and *R*_*n*_ = *nr*_*n*_, so that 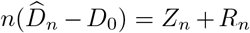. Substitution into (5) yields

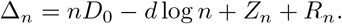

The comparison function *D*(*P*) and its population value *D*_0_ specify the setting targeted by calibration: *D*_0_ > 0 indicates a population-level fitting advantage for the complex model, whereas *D*_0_ ≤ 0 indicates no such advantage. In the latter case, our objective is to construct a threshold that limits the probability of selecting the complex model. The score difference with the remainder removed then satisfies

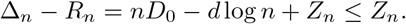

When *D*_0_ ≤ 0, we seek a cutoff *c* ≥ 0 such that

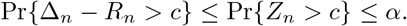

To determine the selection cutoff, we use Cantelli’s inequality to bound the probability that *Z*_*n*_ exceeds this cutoff in terms of its variance. For any random variable *Y* with finite variance *σ*^2^ = Var(*Y*), it gives 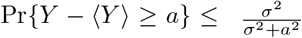 for *a* > 0 [14]. Here, the zero-mean property of *L*_*n*_ implies 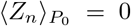. For *Y* = *Z*_*n*_ with 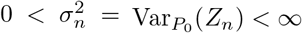, setting *a* = *tσ*_*n*_ gives

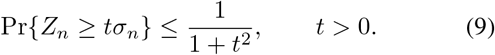

Setting 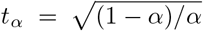 makes the upper bound equal to *α*. This motivates an additional selection margin *t*_*α*_*σ*_*n*_ based on the target level and sampling uncertainty, without changing the BIC penalty. To obtain a computable threshold, we next derive a representation of *Z*_*n*_ from which the unknown scale *σ*_*n*_ can be estimated.

### B. Influence-Function Representation of Sampling Fluctuations

To make the threshold computable, we express the leading random term *Z*_*n*_ as a sum of cell-specific contributions. Let *ψ*(*x*) denote the contribution of a cell with count vector *x* to the first-order sampling fluctuation of the fitted comparison. We seek a representation of the form

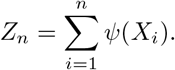

The following conditions support this representation and the subsequent variance estimation.

*Assumption 1(Regularity conditions):*

1. The cell observations are independent and identically distributed under a fixed *P*_0_. The PGF grid is finite and fixed, and *λ* and *α* remain fixed.
2. The matrix Σ(*P*_0_)+*λI*_*m*_ is positive definite. For each model, the population optimum *θ*_0*r*_ is stable, unique and interior. The model PGF is twice continuously differentiable near *θ*_0*r*_, and the Hessian of the fitting criterion at *θ*_0*r*_ is positive definite.
3. The fitted parameters satisfy 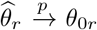, and the numerical optimization errors in the minimized criteria are *o*_*p*_(*n*^−1*/*2^).
4. The contribution function *ψ*(*X*) has finite, strictly positive variance under *P*_0_, and the variance estimator used for calibration converges in probability to this population variance.

Under the model-regularity and fitting conditions in Assumption 1, a first-order sampling expansion [16] yields this representation with 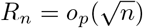. The score decomposition therefore becomes

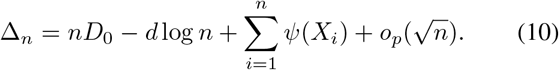

Appendix A, (A.1)–(A.7), derives this expansion from the fitted distance difference.

To use this expansion, we must determine *ψ*(*x*). Resampling changes the empirical PGF, covariance weights and fitted parameters jointly, so each contribution must account for their combined effect on the comparison.

The function *D*(*P*) takes a count distribution *P* as input and returns the difference between two minimized PGF distances; it is therefore a statistical functional. Its influence function measures the first-order response to a small change in probability weights [15]. Under the stated regularity conditions, these local responses also give the observation-level contributions to the sampling expansion [16]. This property allows us to obtain *ψ*(*x*) from the influence function of *D*(*P*).

For a count vector *x*, let *δ*_*x*_ be the point-mass distribution at *x* and let 0 ≤ *ε <* 1 be the perturbation weight. The perturbation path and the influence function at *P*_0_ are

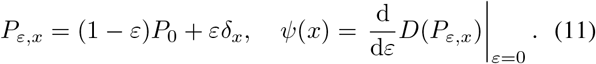

We evaluate this derivative by differentiating each model’s minimum fitting distance in (8). Because both models use the same cells, their responses are calculated along the same perturbation path and then subtracted. At a smooth interior optimum, the parameter gradient vanishes; the envelope property [21] therefore implies that parameter reoptimization contributes no additional first-order term to the minimized distance. We need only differentiate the PGF target and covariance weights, without separately solving for the rate of change of the optimal parameters. This gives

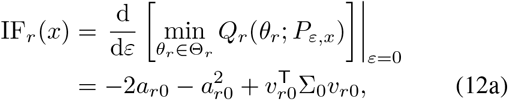

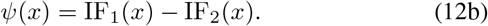

Here, Σ_0_ = Σ(*P*_0_), *W*_0_ = *W*_*λ*_(*P*_0_) and *u*_0_(*x*) = *h*(*x*) − *g*_0_. The remaining quantities and their sample counterparts are

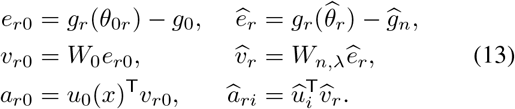

Here, *e*_*r*0_ is the population-optimal PGF residual, *v*_*r*0_ its covariance-weighted form, and *a*_*r*0_ the inner product of *v*_*r*0_ with *u*_0_(*x*); the hatted quantities are their sample counterparts. The dependence of *a*_*r*0_ on *x* is suppressed for brevity. The model-specific influence function IF_*r*_(*x*) measures the sensitivity of the minimum distance, not the distance itself: a positive value indicates an increase to first order, and a negative value the reverse.

Appendix B details the derivation of (12a). Equations (B.1)–(B.2) show why the parameter term vanishes, and (B.16) gives the final expression.

With *ψ*(*x*) specified, we can determine the sampling scale from the variance of a single-cell contribution. Write 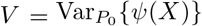. Independence of the cells gives

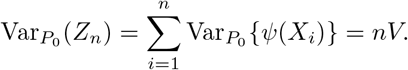

Hence, the scale required by the threshold is 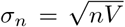. This describes the leading term *Z*_*n*_; the full score difference also contains *R*_*n*_. A computable threshold therefore requires an estimate of *V* from the observed cells and existing model fits.

### C. Data-Driven Threshold and Model Selection

We estimate *V* by first evaluating the influence contributions using the fitted models. At *x* = *X*_*i*_, the quantities *a*_*r*0_, *v*_*r*0_ and Σ_0_ in the closed-form expression (12a) are estimated by *â*_*ri*_, 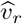 and 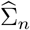, respectively, as defined in (13) and (3). Taking the difference between the two model-specific values for the same cell, as in (12b), yields

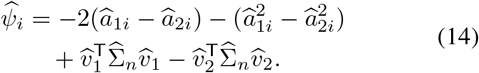

Their centered sample variance provides an estimate of *V* :

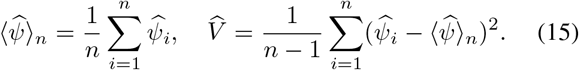

Although the population influence function has zero mean, its estimated values need not average to zero under the covariance convention used here. We therefore subtract their sample mean; the exact centering relation is given in Appendix C, (C.2).

The distinction between the distance and score scales follows from Appendix A, (A.5): the first-order fluctuation of *D*_*n*_ is the average of the independent contributions *ψ*(*X*_*i*_). Its variance is therefore *V/n*, estimated by 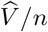. By contrast, *Z*_*n*_ is their sum, so its variance is estimated by 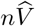.

Thus, 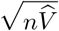 estimates the standard deviation *σ*_*n*_ required by the threshold. Combining it with the multiplier *t*_*α*_ obtained from (9) gives

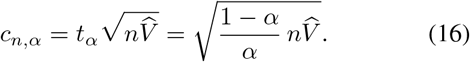

The rule selects *M*_2_ only when Δ_*n*_ > *c*_*n,α*_ and otherwise retains *M*_1_. The complex model’s penalized advantage must therefore exceed a margin set by the estimated sampling uncertainty and the chosen level *α*. For fixed fits, greater estimated uncertainty or a smaller *α* raises this margin. A higher threshold can reduce incorrect selection of the complex model, but may also leave a small, genuine fitting advantage undetected.

This Cantelli-motivated threshold changes only the selection decision. Its computation reuses the fitted residuals and covariance matrix factorization, preserving the original parameter estimates and BIC penalty without resampling or additional optimization. The next section describes the implementation and uses simulations with identical model fits to evaluate how calibration affects model selection.

## IV. Computational Implementation AND Numerical Evaluation

Section III converted the one-sided threshold principle into a computable rule through an influence-function variance estimate. Here, we describe its implementation using the existing PGF fits. With the candidate model fits held unchanged, we improve model-selection accuracy through threshold calibration in the Poisson benchmark introduced in Section II.

### Algorithm 1.

Influence-function-calibrated PGF-BIC

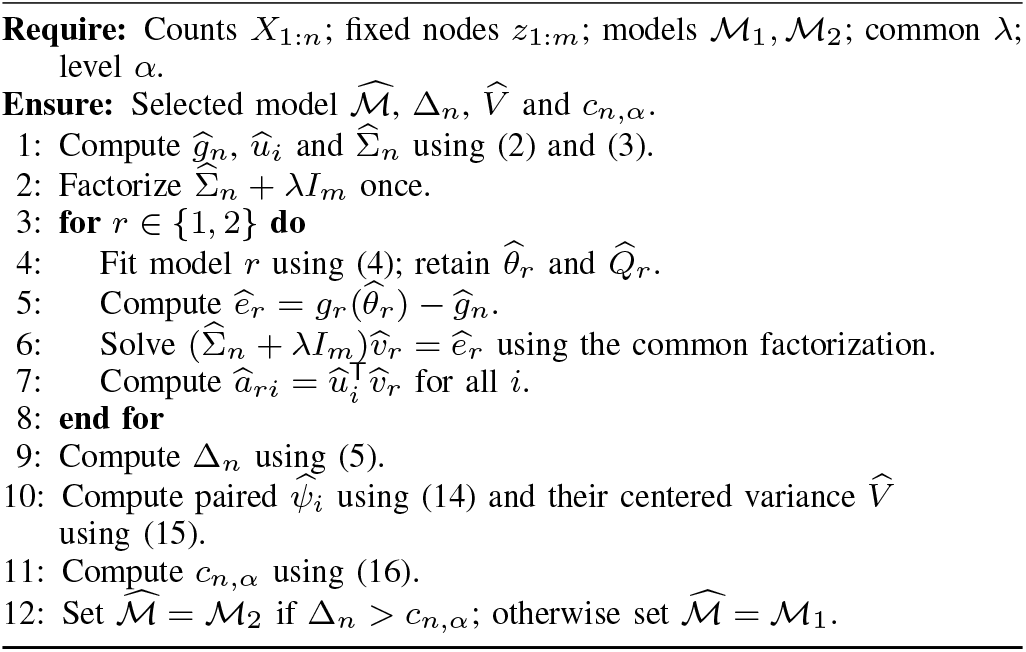

The calibration reuses the existing PGF fits and adjusts only the model-selection threshold, leaving the parameter estimates and fitted distributions unchanged. No resampling or additional optimization is required. Algorithm 1 summarizes the complete procedure.

We apply the procedure to the Poisson(20) datasets introduced in Section II, using the same Poisson and Bursty fits for both selection rules. The grid, ridge parameter, covariance estimates and fitted score differences remain unchanged. With *α* = 0.05, we compare the conventional rule, which selects the Bursty model when Δ_*n*_ > 0, with the calibrated rule, which requires Δ_*n*_ > *c*_*n,α*_. At each of the three sample sizes, performance is measured by the proportion of the 500 independent datasets in which Poisson is correctly selected. Figure 2 compares the correct-selection rates across sample sizes. Calibration improves the correct-selection rate at all three sample sizes: from approximately 30% to approximately 82% at *n* = 100, from 54.6% to 92.4% at *n* = 500, and from 65.4% to 95.4% at *n* = 1000. These results show that threshold calibration reduces incorrect selection of the Bursty model under Poisson-generated data without changing the model fits.

**Fig. 2.**
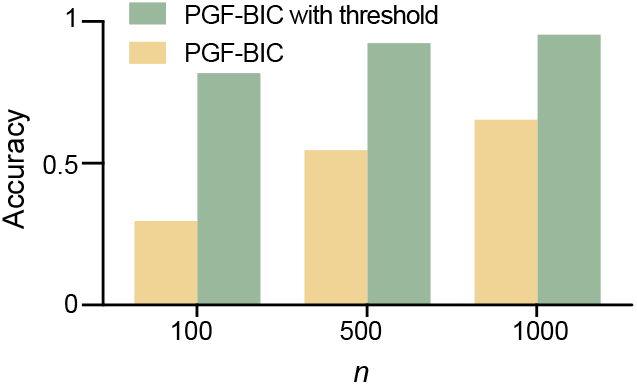
Correct-selection rates for Poisson before and after PGF-BIC threshold calibration (*α*= 0.05) at three sample sizes.

## V. Conclusion

We developed an uncertainty-aware calibration of PGF-BIC [13] that uses influence functions to estimate the sampling variance of the score difference and constructs a selection threshold motivated by Cantelli’s inequality. The Poisson versus Bursty benchmark showed improved correct-selection rates under Poisson-generated data across all three sample sizes. The method preserves the original fits and complexity penalty without resampling or additional optimization, incorporating sampling uncertainty into model selection while maintaining computational efficiency. Further work should extend the first-order theory to boundary and overlapping-model settings and assess the trade-off between reducing incorrect selection of the complex model and detecting its genuine fitting advantages.

## Artificial Intelligence Use

During the preparation of this work the authors used ChatGPT in order to improve the clarity and readability of the manuscript. After using this tool/service, the authors reviewed and edited the content as needed and take full responsibility for the content of the published article.

## AppendixA Derivation OF THE First-Order Expansion OF THE PGF-BIC Difference

This appendix derives (10) directly from the fitted distance difference 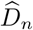, using the influence function already defined in (11).

### 1) Identifying the quantity to expand

Since 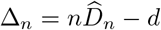 log *n* by (5),

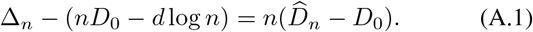

For fixed *n* and model dimensions, the penalty is deterministic. It therefore suffices to expand 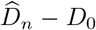 and multiply the result by *n*.

### 2) Expansion in the PGF moment deviations

On the fixed finite PGF grid, the minimum fitting distances depend on the count distribution through the PGF vector *g*_0_ and the second-moment matrix *M*_0_ defined in Section III-A. These determine the covariance through 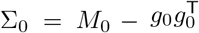. Their empirical counterparts are *ĝ*in (2) and 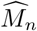, respectively. The expansion is therefore in the deviations *ĝ*_*n*_−*g*_0_ and 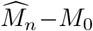, around the population values (*g*_0_, *M*_0_).

Under the smoothness and stable interior-optimum conditions in Assumption 1, the difference of the minimized distances in (8) is locally differentiable in these moment inputs. Let *A*_*j*_ and *B*_*jk*_ denote its partial derivatives with respect to *g*_*j*_ and the distinct second-moment coordinate *M*_*jk*_, respectively, evaluated at (*g*_0_, *M*_0_). We use *j ≤ k* to count each entry of the symmetric matrix only once. These coefficients describe the sensitivity after model optimization and are fixed for a given *P*_0_. Each component of the empirical PGF and raw second-moment matrix is an average of bounded features from independent, identically distributed cells. Its variance is therefore proportional to 1*/n*, and its deviation from the corresponding population moment is *O*_*p*_(*n*^−1*/*2^). A first-order Taylor expansion then gives [16, Thm. 3.1 and its proof, p. 26]

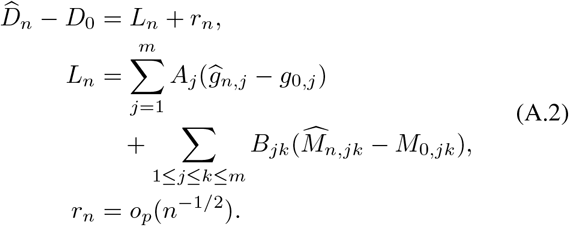

Thus, *L*_*n*_ is the part linear in the moment-estimation errors. The remainder includes the higher-order terms and the numerical optimization error in Assumption 1. The covariance in (3) uses denominator *n −* 1 rather than *n*. Under the stated regularity conditions, this difference leaves the first-order term *L*_*n*_ unchanged and is absorbed into the remainder *r*_*n*_.

### 3) Why the linear term has zero mean

Both moment estimates are averages of fixed functions of the observations. Because every *X*_*i*_ has distribution *P*_0_, linearity of expectation gives

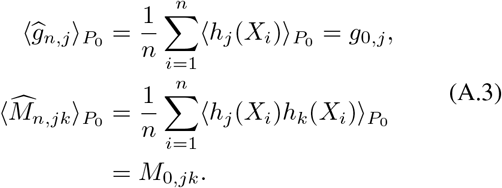

Each moment-estimation error therefore has zero mean. Since *A*_*j*_ and *B*_*jk*_ are evaluated at the population values rather than estimated from the sample, they can be taken outside the expectation:

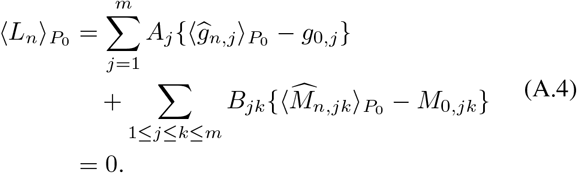

This argument uses linearity of expectation and does not require the moment estimates to be mutually independent. It establishes the zero mean of *L*_*n*_ across repeated samples, not that *L*_*n*_ vanishes in every dataset.

### 4) Relating the linear term to the influence function

Under the perturbation path in (11), the first-order changes in the moment inputs are *h*_*j*_(*x*) − *g*_0,*j*_ and *h*_*j*_(*x*)*h*_*k*_(*x*) − *M*_0,*jk*_. By the chain rule, the same coefficients *A*_*j*_ and *B*_*jk*_ combine these changes into the influence function *ψ*(*x*) already defined there. Averaging over the observations gives 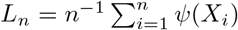, so

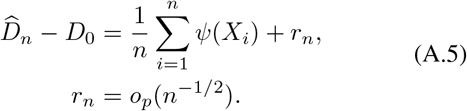

### 5) Recovering the score expansion

Substituting (A.5) into (A.1) gives

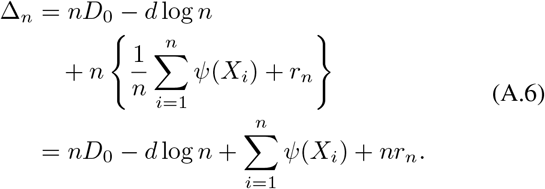

Thus, the decomposition in Section III has 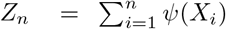 and *R*_*n*_ = *nr*_*n*_. Since *r*_*n*_ = *o*_*p*_(*n*^−1*/*2^), we have 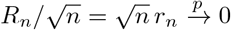, or 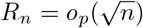. Hence,

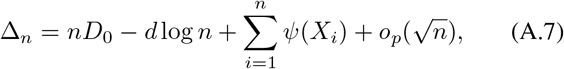

which is (10).

## AppendixB Step-BY-Step Derivation OF THE Model-Specific Influence Function

We start from the derivative of the minimized fitting distance in (12a). After accounting for the change in the optimal parameters, we evaluate the remaining derivative of the weighted criterion to obtain the explicit influence function.

### 1) Accounting for parameter reoptimization

Fix a count vector *x* and follow *P*_*ε,x*_ in (11), with 0 ≤ *ε <* 1. Derivatives at zero are taken along this path from the right. We keep the PGF grid and *λ* fixed, assume Σ_0_ + *λI*_*m*_ is positive definite, and work with a stable, unique interior optimum for each model. The model PGF is twice continuously differentiable near this optimum, and the local Hessian of the fitting criterion is positive definite.

For model *r*, let *θ*_*r*_(*ε*) denote the minimizer along this path, with *θ*_*r*_(0) = *θ*_0*r*_. The local smoothness and stability conditions, together with the nonsingular Hessian, allow the first-order optimality equations to define this differentiable path. The minimized value is therefore *Q*_*r*_(*θ*_*r*_(*ε*); *P*_*ε,x*_). It depends on *ε* through both the optimal parameters and the reference distribution. Starting from (12a), the chain rule gives

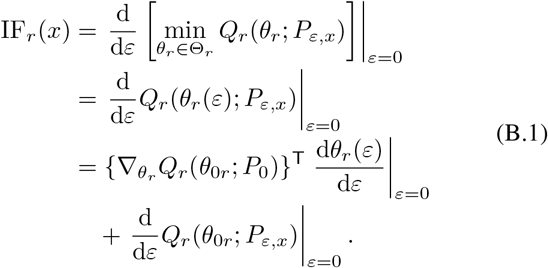

The first term on the last two lines is the contribution from the changing optimal parameters. In the second term, the parameter argument is held at *θ*_0*r*_ and only the distribution is perturbed. At the differentiable interior minimum, 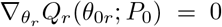, so the coefficient of the parameter derivative vanishes. Hence,

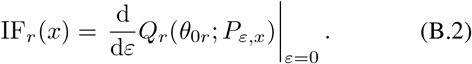

This is the envelope property in the present smooth setting [21]. The effect of parameter reoptimization has been included in (B.1); its first-order contribution is zero at the optimum. Equation (B.2) therefore identifies the derivative that remains to be calculated, without requiring a separate solution for d*θ*_*r*_(*ε*)*/*d*ε*.

### 2) Derivatives required by the weighted criterion

To evaluate (B.2), write the criterion as a quadratic form before differentiating its components. Set *g*_*ε*_ = *g*(*P*_*ε,x*_), Σ_*ε*_ = Σ(*P*_*ε,x*_) and *W*_*ε*_ = *W*_*λ*_(*P*_*ε,x*_). At *ε* = 0, the perturbed distribution coincides with *P*_0_, so *g*_*ε*_ = *g*_0_, Σ_*ε*_ = Σ_0_ and *W*_*ε*_ = *W*_0_. In the following calculations, a prime denotes differentiation with respect to *ε*; for example, 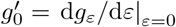. Along this path, write the residual corresponding to (13) as

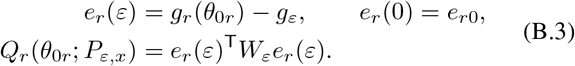

The parameter argument in *e*_*r*_(*ε*) is held at *θ*_0*r*_, so 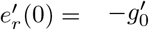. Applying the product rule to the three factors in (B.3) gives

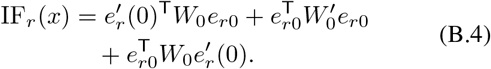

Because *W*_0_ is symmetric, the first and third terms in (B.4) are equal scalars. Combining them and using 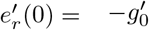 yields

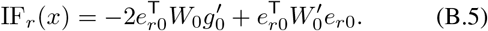

Equation (B.5) identifies the two remaining quantities: 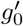 and 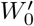. Since *W*_*ε*_ = (Σ_*ε*_+*λI*_*m*_)^−1^, finding 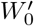 also requires 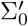. We calculate these derivatives in turn and then substitute them into (B.5).

### 3) PGF derivative

The PGF target is an expectation of the fixed feature vector *h*(*X*). Since *δ*_*x*_ puts all probability at *x*, integration against the mixture is a weighted sum of the two component integrals. For any fixed integrable function *f* with finite *f* (*x*),

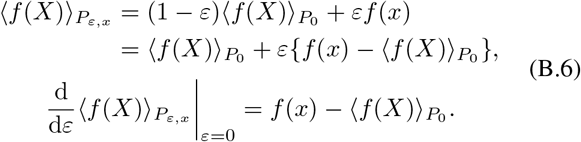

Applying (B.6) to each component of *h*(*X*), and using *u*_0_(*x*) = *h*(*x*) − *g*_0_, gives

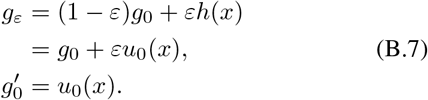

This determines the PGF derivative in (B.5). To obtain the weight derivative, we next calculate how the covariance changes.

### 4) Covariance derivative

The covariance contains both a raw second moment and the product of the mean vector with its transpose. Both terms change under the perturbation. Let 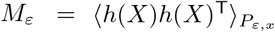. By the definition of covariance, 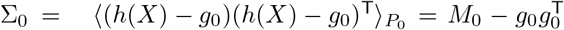, which gives 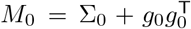. Applying (B.6) entrywise to this fixed matrix-valued function gives

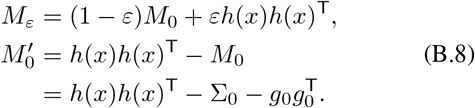

To make the cancellation in the covariance derivative explicit, first expand *h*(*x*) = *g*_0_ + *u*_0_(*x*):

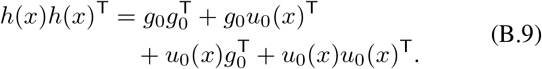

Now differentiate 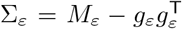 by the product rule. Substituting (B.7) and (B.8), and then cancelling the terms displayed in (B.9), yields

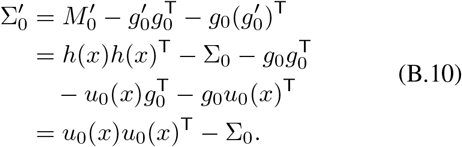

### 5) Weight derivative

The weight matrix is *W*_*ε*_ = (Σ_*ε*_ + *λI*_*m*_)^−1^. To obtain its derivative, start from the defining inverse identity. Since *λ* is fixed, the product rule gives

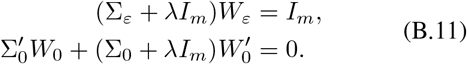

Multiplying the second line from the left by *W*_0_, using *W*_0_(Σ_0_ + *λI*_*m*_) = *I*_*m*_, and moving the first term to the other side gives

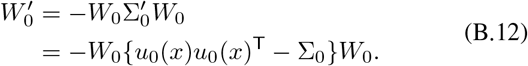

The second equality substitutes (B.10). Both derivatives required by (B.5) are now available, so we can complete the calculation of (B.2).

### 6) Substitution and simplification

First, insert the PGF derivative from (B.7) into (B.5):

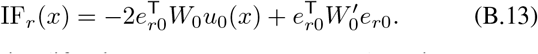

We simplify these two terms separately using *v*_*r*0_ = *W*_0_*e*_*r*0_ and *a*_*r*0_ = *u*_0_(*x*)^T^*v*_*r*0_ from (13). By the symmetry of *W*_0_, the first term becomes

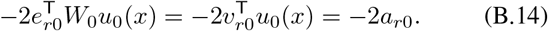

For the second term, substitute the weight derivative from (B.12). Since 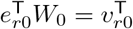

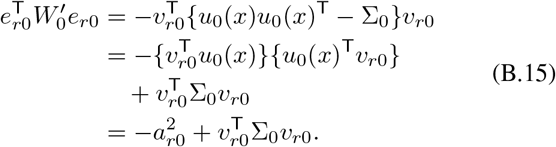

Substituting (B.14) and (B.15) into (B.13) yields

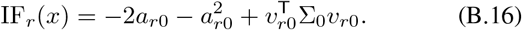

Thus, evaluating the envelope derivative in (B.2) gives the explicit model-specific influence function and completes the derivation of (12a).

## AppendixC Sample Centering OF Estimated Influence Values

This appendix derives the sample mean of the estimated influence values and explains the centering used in (15).

The sample calculation uses 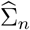 from (3), with denominator *n −* 1, whereas ⟨·⟩_*n*_ averages over *n* observations. The identities ⟨*û*⟩_*n*_ = 0 and 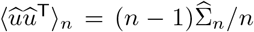, together with the projections in (13), give

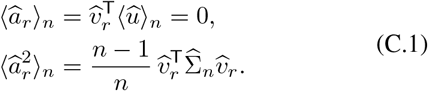

Averaging (14) therefore yields the exact identity

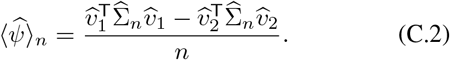

This mean need not be zero. Accordingly, the variance estimator in (15) explicitly subtracts the sample mean before dividing the centered sum of squares by *n* − 1.

