## Supplementary Information for "Uncertainty-Aware Model Selection with a Calibrated Probability-Generating-Function-Based Bayesian Information Criterion"

### Appendix A
### Derivation of the First-Order Expansion of the PGF-BIC Difference

This appendix derives (10) directly from the fitted distance difference $\widehat{D}_n$, using the influence function already defined in (11).

#### 1) Identifying the quantity to expand

Since $\Delta_n = n\widehat{D}_n - d\log n$ by (5),

$$\Delta_n - (nD_0 - d\log n) = n(\widehat{D}_n - D_0). \tag{A.1}$$

For fixed $n$ and model dimensions, the penalty is deterministic. It therefore suffices to expand $\widehat{D}_n - D_0$ and multiply the result by $n$.

#### 2) Expansion in the PGF moment deviations

On the fixed finite PGF grid, the minimum fitting distances depend on the count distribution through the PGF vector $g_0$ and the second-moment matrix $M_0$ defined in Section III-A. These determine the covariance through $\Sigma_0 = M_0 - g_0 g_0^{\mathsf{T}}$. Their empirical counterparts are $\widehat{g}_n$ in (2) and $\widehat{M}_n$, respectively. The expansion is therefore in the deviations $\widehat{g}_n - g_0$ and $\widehat{M}_n - M_0$, around the population values $(g_0, M_0)$.

Under the smoothness and stable interior-optimum conditions in Assumption 1, the difference of the minimized distances in (8) is locally differentiable in these moment inputs. Let $A_j$ and $B_{jk}$ denote its partial derivatives with respect to $g_j$ and the distinct second-moment coordinate $M_{jk}$, respectively, evaluated at $(g_0, M_0)$. We use $j \le k$ to count each entry of the symmetric matrix only once. These coefficients describe the sensitivity after model optimization and are fixed for a given $P_0$. Each component of the empirical PGF and raw second-moment matrix is an average of bounded features from independent, identically distributed cells. Its variance is therefore proportional to $1/n$, and its deviation from the corresponding population moment is $O_p(n^{-1/2})$. A first-order Taylor expansion then gives [16, Thm. 3.1 and its proof, p. 26]

$$\begin{aligned} \widehat{D}_n - D_0 &= L_n + r_n, \\ L_n &= \sum_{j=1}^{m} A_j(\widehat{g}_{n,j} - g_{0,j}) \\ &\quad + \sum_{1\le j\le k\le m} B_{jk}(\widehat{M}_{n,jk} - M_{0,jk}), \\ r_n &= o_p(n^{-1/2}). \end{aligned} \tag{A.2}$$

Thus, $L_n$ is the part linear in the moment-estimation errors. The remainder includes the higher-order terms and the numerical optimization error in Assumption 1. The covariance in (3) uses denominator $n-1$ rather than $n$. Under the stated regularity conditions, this difference leaves the first-order term $L_n$ unchanged and is absorbed into the remainder $r_n$.

#### 3) Why the linear term has zero mean

Both moment estimates are averages of fixed functions of the observations. Because every $X_i$ has distribution $P_0$, linearity of expectation gives

$$\begin{aligned} \langle \widehat{g}_{n,j} \rangle_{P_0} &= \frac{1}{n}\sum_{i=1}^{n} \langle h_j(X_i) \rangle_{P_0} = g_{0,j}, \\ \langle \widehat{M}_{n,jk} \rangle_{P_0} &= \frac{1}{n}\sum_{i=1}^{n} \langle h_j(X_i) h_k(X_i) \rangle_{P_0} \\ &= M_{0,jk}. \end{aligned} \tag{A.3}$$

Each moment-estimation error therefore has zero mean. Since $A_j$ and $B_{jk}$ are evaluated at the population values rather than estimated from the sample, they can be taken outside the expectation:

$$\begin{aligned} \langle L_n \rangle_{P_0} &= \sum_{j=1}^{m} A_j \{\langle \widehat{g}_{n,j} \rangle_{P_0} - g_{0,j}\} \\ &\quad + \sum_{1\le j\le k\le m} B_{jk} \{\langle \widehat{M}_{n,jk} \rangle_{P_0} - M_{0,jk}\} \\ &= 0. \end{aligned} \tag{A.4}$$

This argument uses linearity of expectation and does not require the moment estimates to be mutually independent. It establishes the zero mean of $L_n$ across repeated samples, not that $L_n$ vanishes in every dataset.

#### 4) Relating the linear term to the influence function

Under the perturbation path in (11), the first-order changes in the moment inputs are $h_j(x) - g_{0,j}$ and $h_j(x)h_k(x) - M_{0,jk}$. By the chain rule, the same coefficients $A_j$ and $B_{jk}$ combine these changes into the influence function $\psi(x)$ already defined there. Averaging over the observations gives $L_n = n^{-1}\sum_{i=1}^{n} \psi(X_i)$, so

$$\begin{aligned} \widehat{D}_n - D_0 &= \frac{1}{n}\sum_{i=1}^{n} \psi(X_i) + r_n, \\ r_n &= o_p(n^{-1/2}). \end{aligned} \tag{A.5}$$

#### 5) Recovering the score expansion

Substituting (A.5) into (A.1) gives

$$\begin{aligned} \Delta_n &= nD_0 - d\log n \\ &\quad + n\left\{\frac{1}{n}\sum_{i=1}^{n} \psi(X_i) + r_n\right\} \\ &= nD_0 - d\log n + \sum_{i=1}^{n} \psi(X_i) + nr_n. \end{aligned} \tag{A.6}$$

Thus, the decomposition in Section III has $Z_n = \sum_{i=1}^{n} \psi(X_i)$ and $R_n = nr_n$. Since $r_n = o_p(n^{-1/2})$, we have $R_n/\sqrt{n} = \sqrt{n}\, r_n \xrightarrow{p} 0$, or $R_n = o_p(\sqrt{n})$. Hence,

$$\Delta_n = nD_0 - d\log n + \sum_{i=1}^{n} \psi(X_i) + o_p(\sqrt{n}), \tag{A.7}$$

which is (10).

### Appendix B
### Step-by-Step Derivation of the Model-Specific Influence Function

We start from the derivative of the minimized fitting distance in (12a). After accounting for the change in the optimal parameters, we evaluate the remaining derivative of the weighted criterion to obtain the explicit influence function.

#### *1) Accounting for parameter reoptimization*

Fix a count vector $x$ and follow $P_{\varepsilon,x}$ in (11), with $0 \le \varepsilon < 1$. Derivatives at zero are taken along this path from the right. We keep the PGF grid and $\lambda$ fixed, assume $\Sigma_0 + \lambda I_m$ is positive definite, and work with a stable, unique interior optimum for each model. The model PGF is twice continuously differentiable near this optimum, and the local Hessian of the fitting criterion is positive definite.

For model $r$, let $\theta_r(\varepsilon)$ denote the minimizer along this path, with $\theta_r(0) = \theta_{0r}$. The local smoothness and stability conditions, together with the nonsingular Hessian, allow the first-order optimality equations to define this differentiable path. The minimized value is therefore $Q_r(\theta_r(\varepsilon); P_{\varepsilon,x})$. It depends on $\varepsilon$ through both the optimal parameters and the reference distribution. Starting from (12a), the chain rule gives

$$\begin{aligned}
\mathrm{IF}_r(x) &= \frac{\mathrm{d}}{\mathrm{d}\varepsilon}\left[\min_{\theta_r \in \Theta_r} Q_r(\theta_r; P_{\varepsilon,x})\right]\Bigg|_{\varepsilon=0} \\
&= \frac{\mathrm{d}}{\mathrm{d}\varepsilon} Q_r(\theta_r(\varepsilon); P_{\varepsilon,x})\Bigg|_{\varepsilon=0} \\
&= \{\nabla_{\theta_r} Q_r(\theta_{0r}; P_0)\}^{\mathsf{T}} \frac{\mathrm{d}\theta_r(\varepsilon)}{\mathrm{d}\varepsilon}\Bigg|_{\varepsilon=0} \\
&\quad + \frac{\mathrm{d}}{\mathrm{d}\varepsilon} Q_r(\theta_{0r}; P_{\varepsilon,x})\Bigg|_{\varepsilon=0}.
\end{aligned} \tag{B.1}$$

The first term on the last two lines is the contribution from the changing optimal parameters. In the second term, the parameter argument is held at $\theta_{0r}$ and only the distribution is perturbed. At the differentiable interior minimum, $\nabla_{\theta_r} Q_r(\theta_{0r}; P_0) = 0$, so the coefficient of the parameter derivative vanishes. Hence,

$$\mathrm{IF}_r(x) = \frac{\mathrm{d}}{\mathrm{d}\varepsilon} Q_r(\theta_{0r}; P_{\varepsilon,x})\Bigg|_{\varepsilon=0}. \tag{B.2}$$

This is the envelope property in the present smooth setting [21]. The effect of parameter reoptimization has been included in (B.1); its first-order contribution is zero at the optimum. Equation (B.2) therefore identifies the derivative that remains to be calculated, without requiring a separate solution for $\mathrm{d}\theta_r(\varepsilon)/\mathrm{d}\varepsilon$.

#### *2) Derivatives required by the weighted criterion*

To evaluate (B.2), write the criterion as a quadratic form before differentiating its components. Set $g_\varepsilon = g(P_{\varepsilon,x})$, $\Sigma_\varepsilon = \Sigma(P_{\varepsilon,x})$ and $W_\varepsilon = W_\lambda(P_{\varepsilon,x})$. At $\varepsilon = 0$, the perturbed distribution coincides with $P_0$, so $g_\varepsilon = g_0$, $\Sigma_\varepsilon = \Sigma_0$ and $W_\varepsilon = W_0$. In the following calculations, a prime denotes differentiation with respect to $\varepsilon$; for example, $g_0' = \mathrm{d}g_\varepsilon/\mathrm{d}\varepsilon|_{\varepsilon=0}$. Along this path, write the residual corresponding to (13) as

$$\begin{aligned}
e_r(\varepsilon) &= g_r(\theta_{0r}) - g_\varepsilon, \qquad e_r(0) = e_{r0}, \\
Q_r(\theta_{0r}; P_{\varepsilon,x}) &= e_r(\varepsilon)^{\mathsf{T}} W_\varepsilon e_r(\varepsilon).
\end{aligned} \tag{B.3}$$

The parameter argument in $e_r(\varepsilon)$ is held at $\theta_{0r}$, so $e_r'(0) = -g_0'$. Applying the product rule to the three factors in (B.3) gives

$$\begin{aligned}
\mathrm{IF}_r(x) &= e_r'(0)^{\mathsf{T}} W_0 e_{r0} + e_{r0}^{\mathsf{T}} W_0' e_{r0} \\
&\quad + e_{r0}^{\mathsf{T}} W_0 e_r'(0).
\end{aligned} \tag{B.4}$$

Because $W_0$ is symmetric, the first and third terms in (B.4) are equal scalars. Combining them and using $e_r'(0) = -g_0'$ yields

$$\mathrm{IF}_r(x) = -2e_{r0}^{\mathsf{T}} W_0 g_0' + e_{r0}^{\mathsf{T}} W_0' e_{r0}. \tag{B.5}$$

Equation (B.5) identifies the two remaining quantities: $g_0'$ and $W_0'$. Since $W_\varepsilon = (\Sigma_\varepsilon + \lambda I_m)^{-1}$, finding $W_0'$ also requires $\Sigma_0'$. We calculate these derivatives in turn and then substitute them into (B.5).

#### *3) PGF derivative*

The PGF target is an expectation of the fixed feature vector $h(X)$. Since $\delta_x$ puts all probability at $x$, integration against the mixture is a weighted sum of the two component integrals. For any fixed integrable function $f$ with finite $f(x)$,

$$\begin{aligned}
\langle f(X)\rangle_{P_{\varepsilon,x}} &= (1-\varepsilon)\langle f(X)\rangle_{P_0} + \varepsilon f(x) \\
&= \langle f(X)\rangle_{P_0} + \varepsilon\{f(x) - \langle f(X)\rangle_{P_0}\}, \\
\frac{\mathrm{d}}{\mathrm{d}\varepsilon}\langle f(X)\rangle_{P_{\varepsilon,x}}\Bigg|_{\varepsilon=0} &= f(x) - \langle f(X)\rangle_{P_0}.
\end{aligned} \tag{B.6}$$

Applying (B.6) to each component of $h(X)$, and using $u_0(x) = h(x) - g_0$, gives

$$\begin{aligned}
g_\varepsilon &= (1-\varepsilon)g_0 + \varepsilon h(x) \\
&= g_0 + \varepsilon u_0(x), \\
g_0' &= u_0(x).
\end{aligned} \tag{B.7}$$

This determines the PGF derivative in (B.5). To obtain the weight derivative, we next calculate how the covariance changes.

#### *4) Covariance derivative*

The covariance contains both a raw second moment and the product of the mean vector with its transpose. Both terms change under the perturbation. Let $M_\varepsilon = \langle h(X)h(X)^{\mathsf{T}}\rangle_{P_{\varepsilon,x}}$. By the definition of covariance, $\Sigma_0 = \langle (h(X)-g_0)(h(X)-g_0)^{\mathsf{T}}\rangle_{P_0} = M_0 - g_0 g_0^{\mathsf{T}}$, which gives $M_0 = \Sigma_0 + g_0 g_0^{\mathsf{T}}$. Applying (B.6) entrywise to this fixed matrix-valued function gives

$$
\begin{aligned}
M_\varepsilon &= (1-\varepsilon)M_0 + \varepsilon h(x)h(x)^\top, \\
M_0' &= h(x)h(x)^\top - M_0 \\
&= h(x)h(x)^\top - \Sigma_0 - g_0 g_0^\top.
\end{aligned} \tag{B.8}
$$

To make the cancellation in the covariance derivative explicit, first expand $h(x) = g_0 + u_0(x)$:

$$
\begin{aligned}
h(x)h(x)^\top &= g_0 g_0^\top + g_0 u_0(x)^\top \\
&\quad + u_0(x) g_0^\top + u_0(x) u_0(x)^\top.
\end{aligned} \tag{B.9}
$$

Now differentiate $\Sigma_\varepsilon = M_\varepsilon - g_\varepsilon g_\varepsilon^\top$ by the product rule. Substituting (B.7) and (B.8), and then cancelling the terms displayed in (B.9), yields

$$
\begin{aligned}
\Sigma_0' &= M_0' - g_0' g_0^\top - g_0 (g_0')^\top \\
&= h(x)h(x)^\top - \Sigma_0 - g_0 g_0^\top \\
&\quad - u_0(x) g_0^\top - g_0 u_0(x)^\top \\
&= u_0(x) u_0(x)^\top - \Sigma_0.
\end{aligned} \tag{B.10}
$$

*5) Weight derivative*

The weight matrix is $W_\varepsilon = (\Sigma_\varepsilon + \lambda I_m)^{-1}$. To obtain its derivative, start from the defining inverse identity. Since $\lambda$ is fixed, the product rule gives

$$
\begin{aligned}
(\Sigma_\varepsilon + \lambda I_m) W_\varepsilon &= I_m, \\
\Sigma_0' W_0 + (\Sigma_0 + \lambda I_m) W_0' &= 0.
\end{aligned} \tag{B.11}
$$

Multiplying the second line from the left by $W_0$, using $W_0(\Sigma_0 + \lambda I_m) = I_m$, and moving the first term to the other side gives

$$
\begin{aligned}
W_0' &= -W_0 \Sigma_0' W_0 \\
&= -W_0 \{u_0(x) u_0(x)^\top - \Sigma_0\} W_0.
\end{aligned} \tag{B.12}
$$

The second equality substitutes (B.10). Both derivatives required by (B.5) are now available, so we can complete the calculation of (B.2).

*6) Substitution and simplification*

First, insert the PGF derivative from (B.7) into (B.5):

$$
\mathrm{IF}_r(x) = -2 e_{r0}^\top W_0 u_0(x) + e_{r0}^\top W_0' e_{r0}. \tag{B.13}
$$

We simplify these two terms separately using $v_{r0} = W_0 e_{r0}$ and $a_{r0} = u_0(x)^\top v_{r0}$ from (13). By the symmetry of $W_0$, the first term becomes

$$
-2 e_{r0}^\top W_0 u_0(x) = -2 v_{r0}^\top u_0(x) = -2 a_{r0}. \tag{B.14}
$$

For the second term, substitute the weight derivative from (B.12). Since $e_{r0}^\top W_0 = v_{r0}^\top$,

$$
\begin{aligned}
e_{r0}^\top W_0' e_{r0} &= -v_{r0}^\top \{u_0(x) u_0(x)^\top - \Sigma_0\} v_{r0} \\
&= -\{v_{r0}^\top u_0(x)\}\{u_0(x)^\top v_{r0}\} \\
&\quad + v_{r0}^\top \Sigma_0 v_{r0} \\
&= -a_{r0}^2 + v_{r0}^\top \Sigma_0 v_{r0}.
\end{aligned} \tag{B.15}
$$

Substituting (B.14) and (B.15) into (B.13) yields

$$
\mathrm{IF}_r(x) = -2 a_{r0} - a_{r0}^2 + v_{r0}^\top \Sigma_0 v_{r0}. \tag{B.16}
$$

Thus, evaluating the envelope derivative in (B.2) gives the explicit model-specific influence function and completes the derivation of (12a).

### Appendix C
### Sample Centering of Estimated Influence Values

This appendix derives the sample mean of the estimated influence values and explains the centering used in (15).

The sample calculation uses $\widehat{\Sigma}_n$ from (3), with denominator $n-1$, whereas $\langle\cdot\rangle_n$ averages over $n$ observations. The identities $\langle \widehat{u} \rangle_n = 0$ and $\langle \widehat{u}\widehat{u}^\top \rangle_n = (n-1)\widehat{\Sigma}_n / n$, together with the projections in (13), give

$$
\begin{aligned}
\langle \widehat{a}_r \rangle_n &= \widehat{v}_r^\top \langle \widehat{u} \rangle_n = 0, \\
\langle \widehat{a}_r^2 \rangle_n &= \frac{n-1}{n} \widehat{v}_r^\top \widehat{\Sigma}_n \widehat{v}_r.
\end{aligned} \tag{C.1}
$$

Averaging (14) therefore yields the exact identity

$$
\langle \widehat{\psi} \rangle_n = \frac{\widehat{v}_1^\top \widehat{\Sigma}_n \widehat{v}_1 - \widehat{v}_2^\top \widehat{\Sigma}_n \widehat{v}_2}{n}. \tag{C.2}
$$

This mean need not be zero. Accordingly, the variance estimator in (15) explicitly subtracts the sample mean before dividing the centered sum of squares by $n-1$.
